# Plant conservation assessment at scale: rapid triage of extinction risks

**DOI:** 10.1101/2022.08.16.503993

**Authors:** Taylor AuBuchon-Elder, Patrick Minx, Bess Bookout, Elizabeth A. Kellogg

**Author notes:** **Correspondence** Taylor AuBuchon-Elder, Donald Danforth Plant Science Center, 975 North Warson Road, St. Louis, MO, USA.

## Abstract

**Summary:**

- The IUCN Red List criteria are widely used to determine extinction risks of plant and animal life. Here, we use The Red List’s criterion B, Geographic Range Size, to provide preliminary conservation assessments of the members of a large tribe of grasses, the Andropogoneae, with ∼1100 species, including maize, sorghum, and sugarcane and their wild relatives.
- We use georeferenced occurrence data from the Botanical Information and Ecology Network (BIEN) and automated individual species assessments using ConR to demonstrate efficacy and accuracy in using time-saving tools for conservation research. We validate our results with those from the IUCN-authorized assessment tool, GeoCAT.
- We discovered a remarkably large gap in digitized information, with slightly more than 50% of the Andropogoneae lacking sufficient information for assessment. ConR and GeoCAT largely agree on which taxa are of least concern (>90%) or possibly threatened (<10%), highlighting that automating assessments with ConR is a viable strategy for preliminary conservation assessments of large plant groups. Results for crop wild relatives are similar to those for the entire data set.
- Increasing digitization and collection needs to be a high priority. Available rapid assessment tools can then be used to identify species that warrant more comprehensive investigation.

**Societal Impact Statement:** The current rate of global biodiversity loss creates a pressing need to increase efficiency and throughput of extinction risk assessments in plants. We must assess as many plant species as possible, working with imperfect knowledge, to address the habitat loss and seemingly countless extinction threats of the Anthropocene. Large-scale, preliminary conservation assessments can play a fundamental role in setting priorities for more in-depth investigation.

## 1 INTRODUCTION

Anthropogenic change has led to steep drops in biodiversity in every biome on the planet (Hautier et al., 2015), contributing to what some suggest is the world’s sixth mass extinction (Kling et al., 2018; Cowie et al., 2022). The essential role of biodiversity in ecosystem function and ecosystem services is well-established (Díaz et al., 2006), and plants are integral players in these services (Pelletier et al., 2018). However, knowledge of the extinction risk of plant species is patchy, leaving us in the dark about rates of ecosystem decline or biodiversity loss (Panter et al., 2020; Rivers et al., 2011), and hindering our ability to mitigate and prioritize risks to wild plant species. Declining diversity among crop wild relatives is also cause for concern (Khoury et al., 2022) because of their importance in agricultural research and food security (Castañeda-Álvarez et al., 2016; Khoury et al., 2020). The Global Strategy for Plant Conservation (GSPC) of the 2011-2020 United Nations Convention on Biological Diversity (CBD) called for assessment of the conservation status of all known plant species, but this ambitious goal remains unattained.

The International Union for Conservation of Nature (IUCN) Red List criteria (Figure 1) provide the authoritative framework for assessing conservation status and contributing to the GSPC objectives (Sharrock, 2020). Criterion B (species geographic range) is most commonly used for plants, as assessments can be done using only georeferenced locality data. The major subcriteria are B1, Extent of Occurrence (EOO), and B2, Area of Occupancy (AOO), where EOO is the area encompassed by the minimum convex or alpha hull that includes all reported locations, and AOO determines the number of grid cells of a specified size within the EOO occupied by the species (Dauby et al., 2017). These values permit determination of a taxon’s conservation status as: 1) Data Deficient (DD), 2) Least Concern (LC), 3) Near Threatened (NT), 4) Vulnerable (VU), 5) Endangered (EN), 6) Critically Endangered (CR), 7) Extinct in the Wild (EW), or 8) Extinct (EX), with varying specificity and subcriteria dependent on the data available (IUCN, 2012).

**Figure 1.**
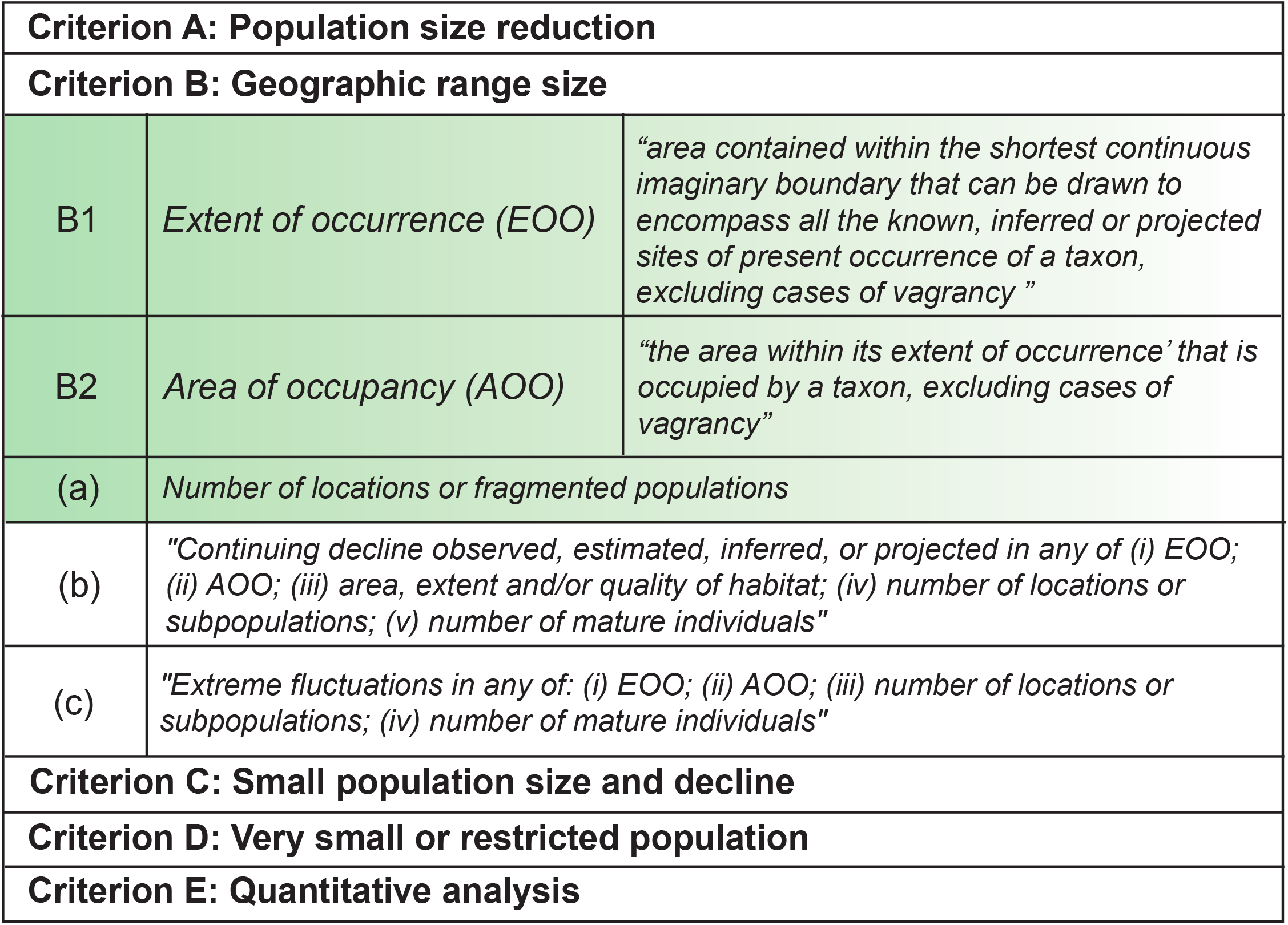
IUCN criteria for assessing risk of extinction, redrawn and directly quoted from IUCN (2012). Criteria with green shading evaluated in this paper. Note that criteria A, C, and B2b and c all require repeated assessment whereas B1, B2, and B2a can be estimated from single records.

While Red List methods are the gold standard for conservation assessment, they can require extensive expertise, time, funding, and geographic accessibility, and for logistical reasons are not always feasible (Rivers et al., 2011; Le Breton et al., 2019). Biases in number of assessments, geographic preferences, and organism types also exacerbate the gaps in conservation data (Walker et al., 2020). Among plants, woody perennials and species with known human use are over-represented in Red List assessments whilst Lamiaceae, Orchidaceae, Poaceae, and Asteraceae are notably under-represented (Lughadha et al., 2020). Only 10.5% of the ∼383,670 species of vascular plants have been globally assessed by the IUCN Red List (Lughadha et al., 2016; Holz et al., 2022; Lughadha et al., 2020), well short of the GSPC 2020 targets.

Documenting and assessing plant species at a speed that matches the urgency of the extinction crisis requires high throughput methods, even if they are imperfect. Such preliminary conservation assessments can then be shared with on-the-ground experts and stakeholders with the critical local knowledge, resources, and community connections to a) confirm or deny the preliminary conservation status conclusion, b) provide expert detail on current threats and population outlooks beyond those available in on-line databases, and c) set priorities for species and/or habitat conservation.

Here, we assess extinction risk for a large clade of plants, the grass tribe Andropogoneae (Poaceae: Panicoideae), using automated tools for download and analysis of species occurrences, and have validated the approach with less automated methods. The tribe includes ca. 1,100 species of grasses that are prevalent in many of the world’s most endangered ecosystems (Estep et al., 2014; Lehmann et al., 2019; Scholtz & Twidwell, 2022). They are the ecologically dominant species in the North American tallgrass prairie, African savannas, and south Asian tropical grasslands (Figure 2). Andropogoneae are often used for forage, aid in erosion mitigation, and may provide bigger carbon sinks than forests (Dass et al., 2018). In addition, the tribe includes some of the world’s most aggressive weeds, including *Sorghum halepense, Imperata cylindrica*, and *Heteropogon contortus*.

**Figure 2.**
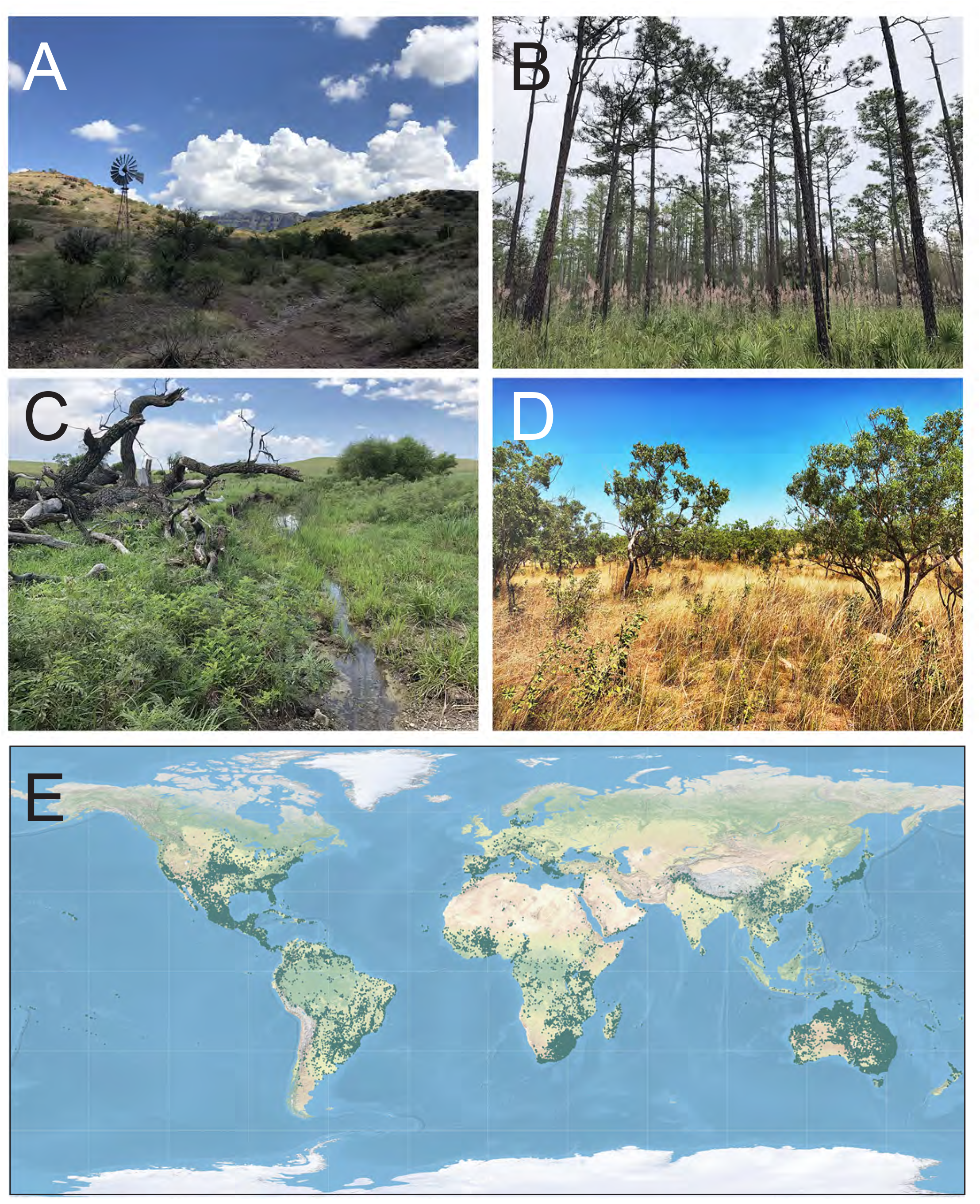
Ecosystems where Andropogoneae naturally occur. Clockwise from top left: (A) Upper riparian sloped grassland in southern Arizona, US; (B) Mesic longleaf pine flatwoods in central Florida, US; (C) Bottomland tallgrass prairie in Flint Hills, Kansas, US; (D) Dry tropical savanna in Northern Territory, Australia. (E) Estimated range map of Andropogoneae (excluding *Zea mays, Sorghum bicolor*, and *Sorghum halepense*) using available locality data (colored in green). Mapped with QGIS-LTR Version 3.22.4. Graticules from Natural Earth. Photos taken by TAE.

Andropogoneae also includes some of the world’s most valuable crop species (*Zea mays, Sorghum bicolor, Saccharum officinarum*) and their wild relatives (*Tripsacum, Miscanthus*). Research on the tribe thus improves cereal crop efficiency and agricultural sustainability due to their highly adaptive C_4_ photosynthesis and drought tolerance (Hattersley and Watson, 1992). In contrast to the well-known crops, the tribe also includes many species about which we know little beyond their original descriptions.

Only 100 species of Andropogoneae have been assessed in the IUCN Red List (Supplemental Table 1), including 26 crop wild relatives plus *Zea mays* itself. We have performed preliminary conservation assessments for the remaining 1100 species, based on the Red List Criteria EOO and AOO. We used one automated tool, ConR, and compared it to the current manual standard, GeoCAT, for validation (Bachman et al., 2011). For georeferenced locality data, we started with the Botanical Information and Ecology Network (BIEN)(Enquist et al., 2016) and added extensive taxonomic curation to account for synonymy and other data artifacts such as misspelling of names. We demonstrate that just under half of the species in the tribe can be assessed under criterion B, with the remainder having few or no records in online databases. Of those that could be assessed, a large majority (91%) appear not to be globally threatened, permitting future studies to prioritize the 46-50 species that may be at risk. Most of the potentially threatened species are severely under-collected and/or digitized, with fewer than 10 unique georeferenced occurrences. This alone may be reason enough to focus first on those species.

## 2 MATERIAL AND METHODS

Our workflow involved: 1) assembling a comprehensive list of species names; 2) taxonomic reconciliation; 3) retrieval and organization of occurrence data, and 4) preliminary conservation assessment of each species (Figure 3). Taxonomic and occurrence data were provided by BIEN (Enquist et al., 2016), accessed and manipulated with the R package BIEN R (Maitner, 2020). Some resources in BIEN are similar to those in the Global Biodiversity Information Facility (GBIF), but BIEN provides additional data cleaning, with species occurrence data validated for spatial errors (Maitner et al., 2018); it also retrieves more occurrence records per species than GBIF (Panter et al., 2020).

**Figure 3.**
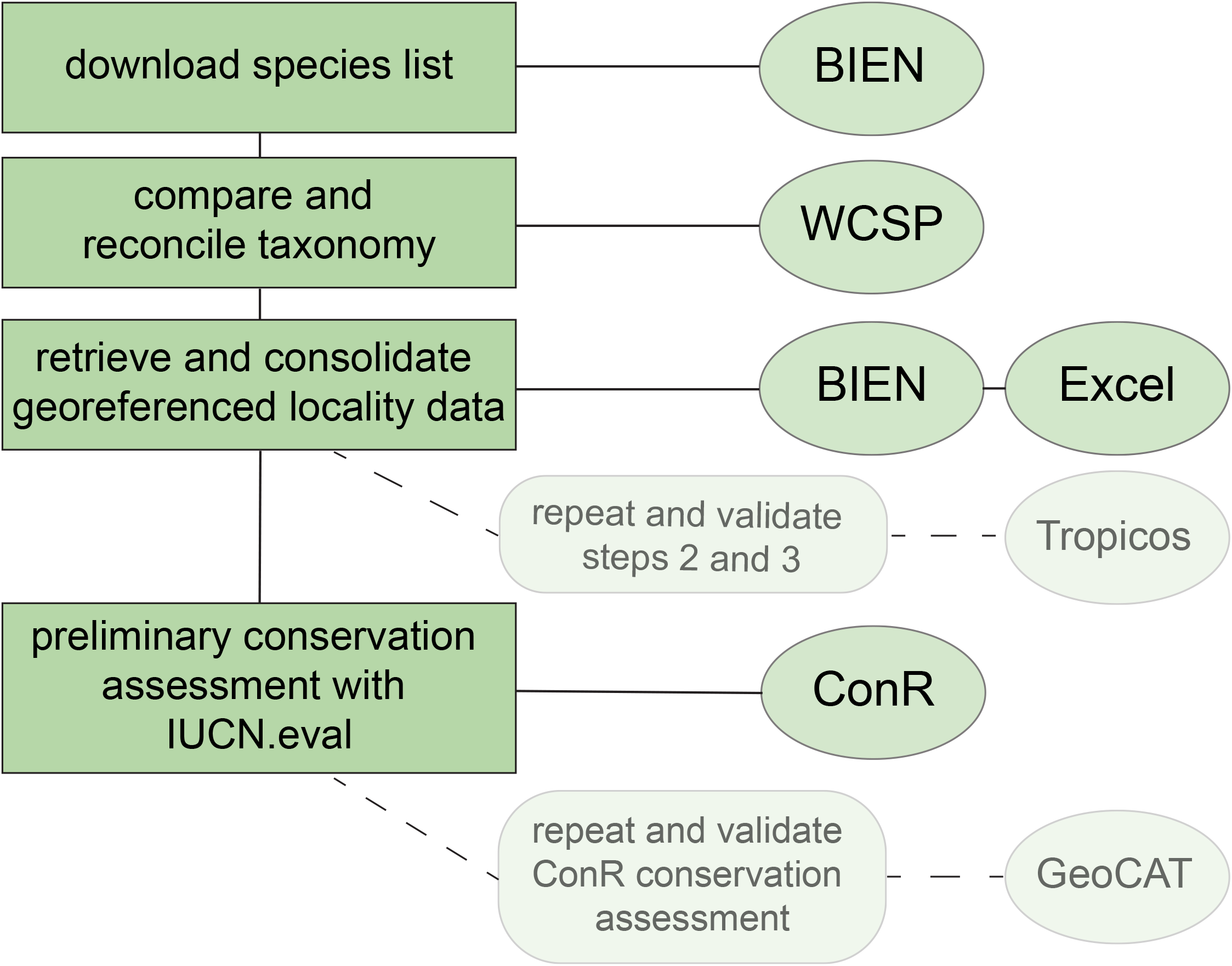
Steps in workflow for conservation assessment of Andropogoneae. Primary workflow in boxes on the left of the figure. Tools used in ovals on the right. Validation steps used for this paper in rounded rectangles with lighter fill in the center.

### 2.1 Databases and comprehensive lists of species names

A list of all Poaceae species in the BIEN database was downloaded using the query [BIEN_full_taxonomy_family], and then filtered to retain only species in genera assigned to Andropogoneae by GrassBase (Clayton, W.D. et al., 2006) and the more recent World Checklist of Selected Plant Families (http://wcsp.science.kew.org/) (hereafter, “Kew”).

Our query retrieved all recorded species names, whether or not they were currently accepted. Many are placed in synonymy by GrassBase, WCSP or BIEN itself via the Taxonomic Names Resolution Service (TNRS) (Boyle et al., 2013), although the most recent release of the TNRS (2021) appeared after the taxonomic work described here was complete. The list also included names that were misspelled, attributed to the wrong taxonomic authority, or given the wrong Latin ending. We wished to capture records associated with as many species names as possible to avoid missing localities digitized under outdated synonyms or erroneous names, which are often attached to only a few records and thus may appear, incorrectly, as threatened. After retrieving all species names in all Andropogoneae genera, we had two species lists: a) the authoritative Kew list (1,157 names) and b) the BIEN list (1,556 names). Lists were compared side by side in a single .csv file.

### 2.2 Taxonomic reconciliation

We compared the two lists using the Excel code (=IFERROR(VLOOKUP(A2,$D$2:$D$1556,1,0),(“not in BIEN”))) or “not in Kew”), where A and D are the columns Kew and BIEN, respectively and 1556 is the number of rows (names). Exact matches were placed into an “Exact match” list. Species names appearing only in the BIEN list were manually checked against WCSP. If the species was accepted in WCSP, we added it into a “consolidated working list” (the “Exact match list” + new additions). If the species was a synonym, it was recorded along with the preferred name. If the preferred accepted name was still valid but not already in the “consolidated working list”, it was added. We also verified the taxonomic authority of each name. Duplicates produced by minor misspellings or discrepancies in gender determinations were manually corrected, using WCSP as the authority. Species names that appeared only in BIEN, did not have synonyms, and appeared as “Unplaced” in the WCSP database were removed.

The “Cleaned Species List” then included: 1) exact matches between the two lists; 2) accepted names only in the Kew list; 3) accepted names only in the BIEN list plus their Kew list synonyms, and 4) unplaced names. This produced a final working data set with 1,130 unique names, a number slightly lower than the number of 1,224 species estimated by Welker et al. (2020).

### 2.3 Retrieval of occurrence data

We downloaded occurrence data for all genera in the Cleaned Species List with [BIEN_occurrence_genus], retrieving information for political boundaries and collection information [political.boundaries=TRUE, collection.info=TRUE]. This returned a dataframe containing all available occurrence records for each genus and species: scrubbed genus and species, country, state, county, locality, latitude, longitude, date collected, datasource, dataset, data owner, data source ID, catalog number, identified by, and date identified. Names without occurrences were automatically skipped by the program. Occurrences for BIEN-only synonyms were consolidated with those of their accepted name.

To validate our query of BIEN, we repeated the same process but with the Tropicos database (https://Tropicos.org). We reconciled names between Tropicos and Kew and then retrieved all available coordinates from Tropicos for each species. While nearly all results were the same, we gained usable localities for nine species that lacked coordinates in BIEN.

Andropogoneae already in IUCN’s Red List assessment data were kept in our analyses (Supplemental Table 2). *Zea mays* and *Sorghum bicolor* were omitted because they are cultivated worldwide; *S. halepense* was omitted because it is an aggressive weed with a global distribution. The final number of species for assessment was 1,127.

### 2.4 Conservation assessments of accepted species

Extinction risk was assessed with the R-based tool ConR (Dauby et al., 2017) and validated with the web-based and IUCN-approved tool GeoCat (Bachman et al., 2011). Both tools automate risk assessment and generation of maps. Comma-separated files (.csv) with species name, latitude, and longitude were extracted from the main dataframes and as specified by each program. ConR uses batch uploads and automatically generates spreadsheets of the results (Dauby et al., 2017). Within ConR, we used the modules [EOO.computing] and [IUCN.eval]. For comparison, we also used GeoCAT with grid cells of 2km by 2km, as recommended (IUCN, 2012). GeoCAT lacks a batch upload option and requires manual recording of outputs.

Both analytical tools produced a) EOO km^2^ and AOO km^2^; b) EOO- and AOO-based risk ratings; and c) number of unique occurrences analyzed. ConR also automatically estimated number of locations, where “location” is “a geographically or ecologically distinct area in which a single threat can rapidly affect all individuals of the taxon present” (IUCN, 2012). ConR uses only unique occurrences and skips exact duplicates, whereas GeoCAT includes the duplicates, but these do not change the convex hulls so EOO and AOO values are unaffected. EOO values for ConR and GeoCAT were compared in bivariate plots to identify outliers, which were manually checked; all represented typographical errors that were corrected.

ConR could not compute EOO for 62 species with ranges spanning the 180th meridian, although GeoCAT could. We manually inspected the GeoCAT maps and found that 15 species included vagrants that could be manually removed and the analysis re-run. ConR analyzed seven of them, whereas the remaining species still had ranges that were too broad.

## 3 RESULTS

### 3.1 Fewer than half of Andropogoneae species had enough georeferenced data for assessment

The reconciled species list included 1,130 unique species of Andropogoneae (Supplemental table 2), retrieved 59,186 individual occurrences (as of 2021); eliminating two crops (*Zea mays, Sorghum bicolor*) and one aggressive weed (*Sorghum halepense*) gave a data set of 1,127 species. The conservation status of 573 (51%) of them could not be rapidly assessed with ConR.

Of the 573, 337 had no accessible georeferenced occurrence data. Of the 337, 18 had already been assessed in the IUCN Red List, so we inferred their localities were protected or did not have digitized specimen data. The other 319 lacked digitized coordinates. An additional 181 species had only 1-2 occurrences per species, which prevented preliminary assessments because EOO estimates require at least three occurrences. The remaining 55 species could be assessed by GeoCat but not ConR because ConR does not provide results for species with ranges that cross the 180th meridian (Supplemental table 2).

Lack of data could reflect low collection numbers, lack of digitization, or locality data without georeferencing, among other causes. The five countries with the most data deficient species were India, Myanmar, China, Thailand, and Vietnam, each with more than 50 species without digitized georeferenced locality data (∼223 species, 78, 62, 78, and 51, respectively) (Figure 4).

**Figure 4.**
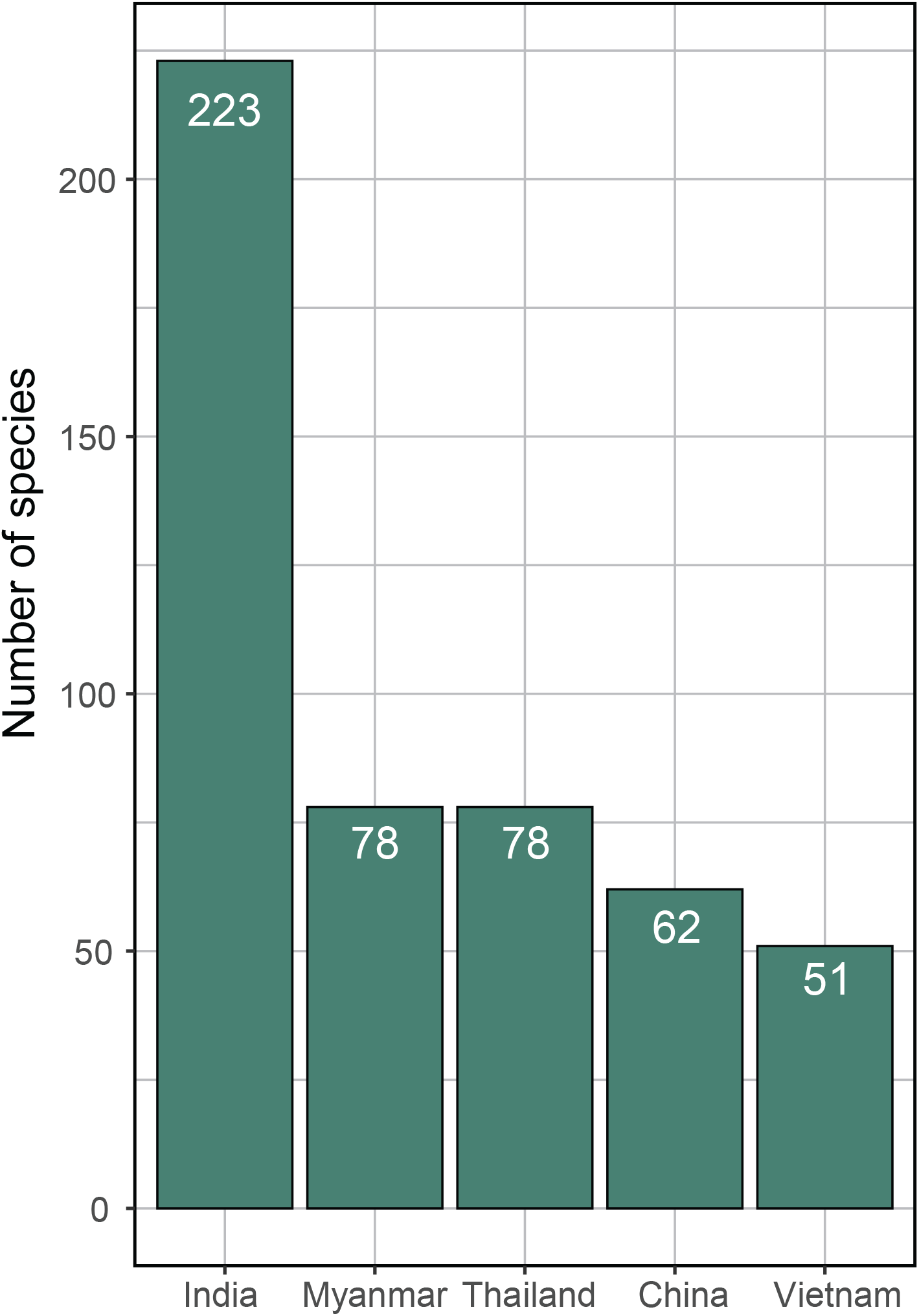
Countries with the largest numbers of species lacking digitized locality data. Number of species based on reported native ranges based on political boundaries (Plants of the World Online (POWO), accessed December 2021).

### 3.2 9% of rapidly assessed Andropogoneae are potentially threatened

554 species (49% of the tribe) could be assessed by both GeoCAT and ConR, which produced EOO and AOO values that were nearly identical (EOO, r^2^=1; AOO, r^2^=0.998). Most of the species (504/554 or 91%) were assessed by both tools as LC or NT. GeoCAT listed 487 of these as LC and 17 as NT, whereas ConR does not distinguish LC from NT. Of the 55 species with coordinates that cross the 180th meridian, GeoCAT assessed all as being LC, and confirmed 10 species previously assessed for the IUCN Red List as LC.

The remaining 9% (50/554) of the assessed species were assessed as CR, EN, or VU by one or both tools. Of these, 46 appeared threatened (2 CR, 26 EN, 18 VU) according to ConR, or 50 (7 CR, 28 EN, 13 VU) according to GeoCAT. ConR did not indicate that *Microstegium japonicum* and *Rottboellia parodiana* were threatened, although GeoCAT did, despite having similar EOO values. Subcategories of potentially threatened status (CR, EN, VU) varied more between the two tools. 26% of the 50 potentially threatened species differed in their EOO letter grade (Table 1).

**Table 1.**
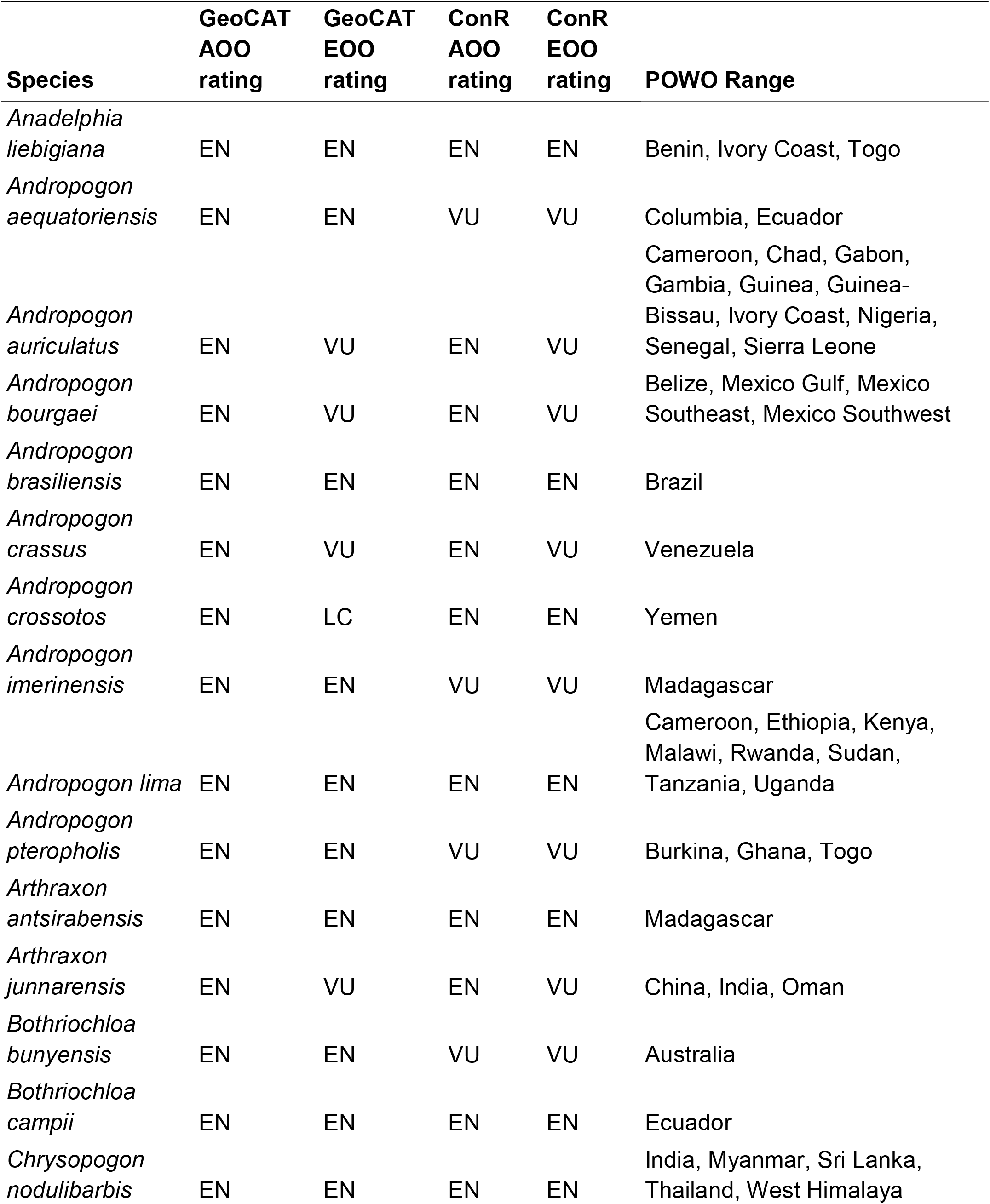
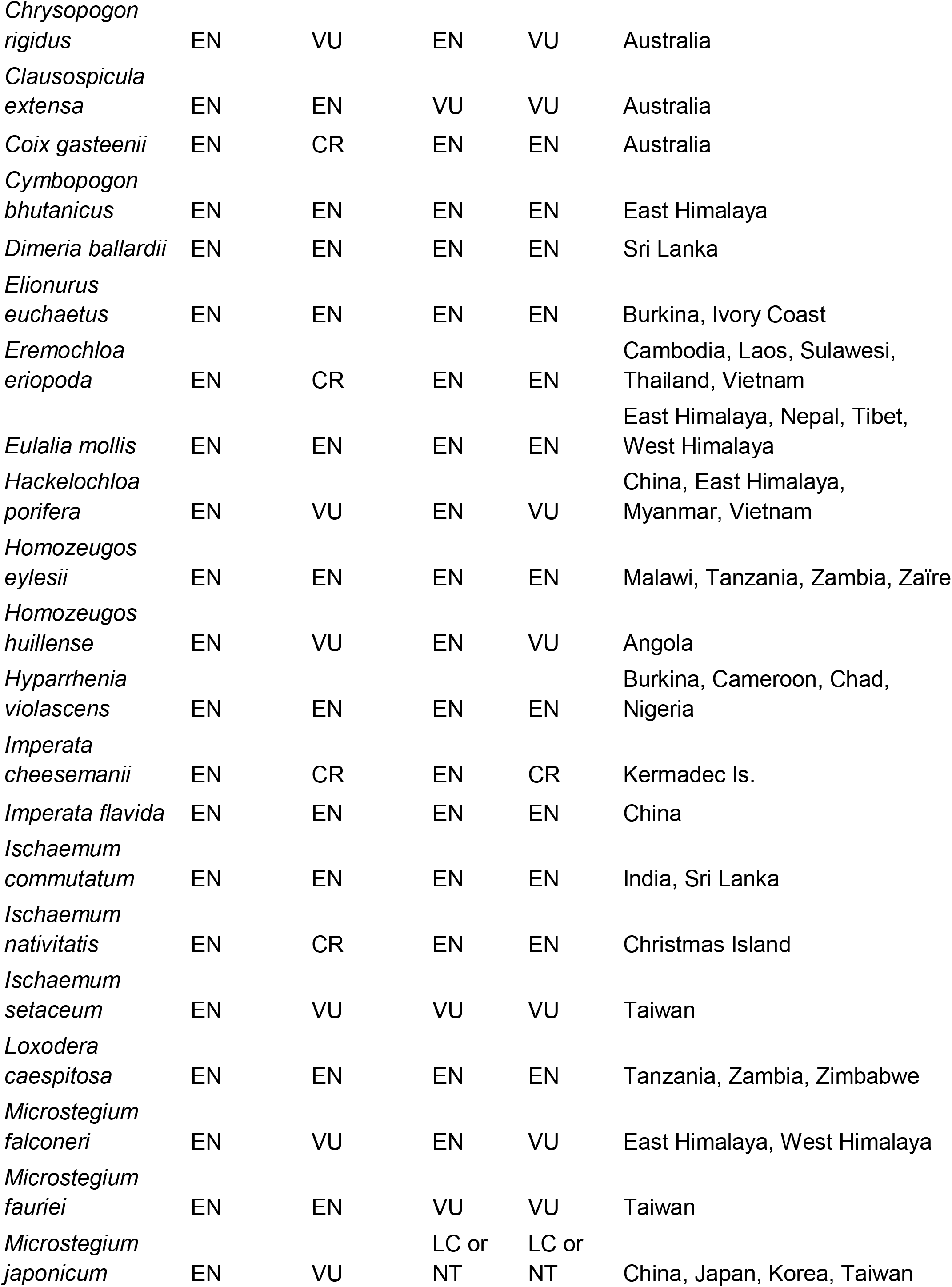
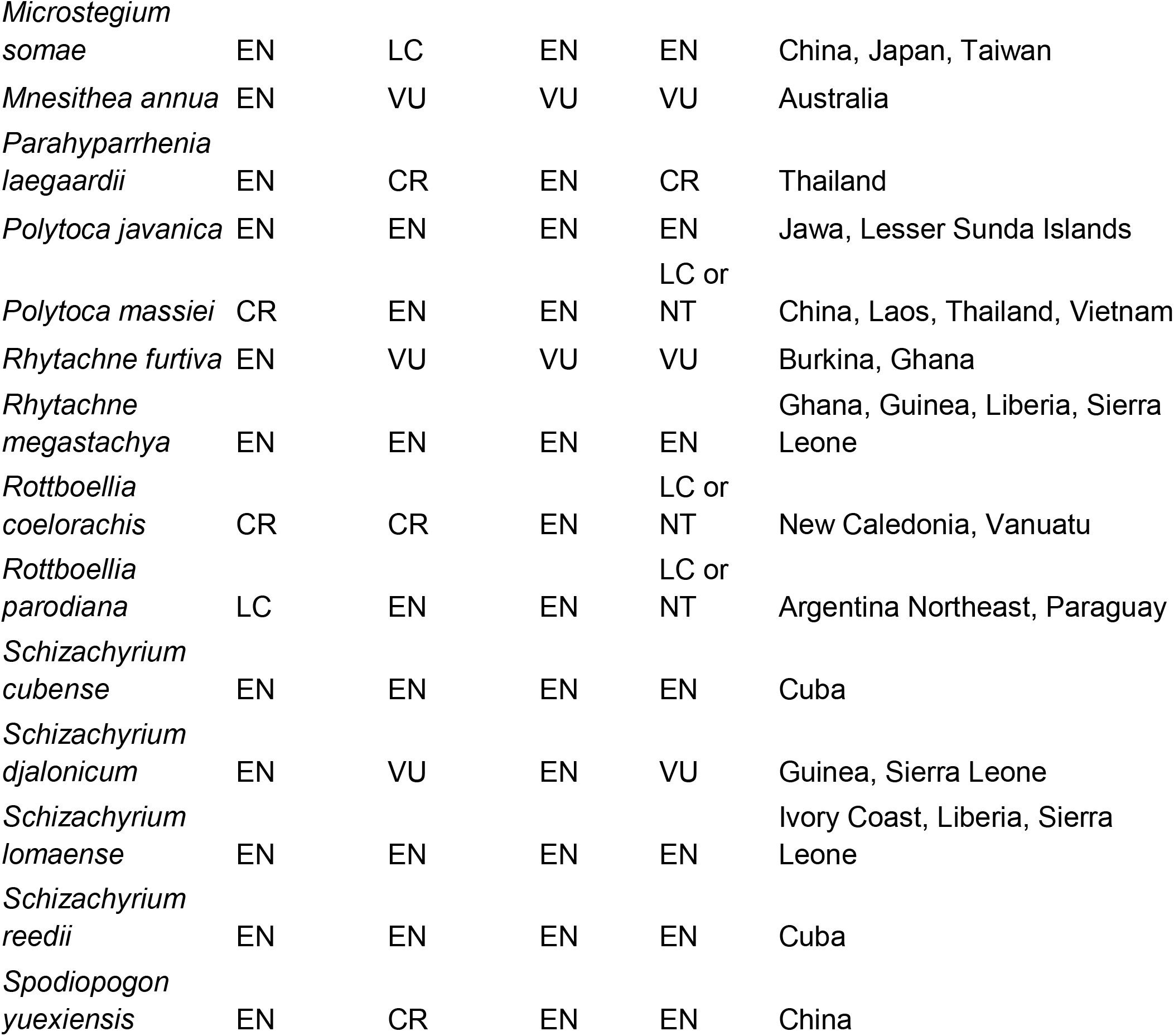
Andropogoneae species identified in this analysis as possibly threatened by either GeoCAT, ConR or both. 46 species appeared threatened in ConR versus 50 in GeoCat. Potentially threatened species are not concentrated in one region. AOO, area of occupancy; EOO, extent of occurrence; POWO, Plants of the World Online.

### 3.3 Phylogenetic clustering

We explored whether potentially threatened species were concentrated in particular clades using the plastome phylogeny and subtribal classification of Andropogoneae provided by Welker et al. (2020) (Figure 5). Each subtribe contained a few (generally 1-8) species that were potentially threatened, although the largest subtribe, Andropogoninae, had 30. To normalize the number of threatened species by size of the subtribe, we used species numbers provided by Welker et al. (2020). The WCSP species list (Supplemental Table 2) does not provide subtribal assignments, and Welker et al. (2020) provide only species numbers, not lists, for their subtribes, estimates of species numbers are approximate. In addition, a few genera listed by WCSP are polyphyletic in the phylogeny. This polyphyly affects the genera *Phacelurus*, one species of which is segregated as the genus *Jardinea*, and *Rottboellia*, some species of which are segregated as *Mnesithea* by Welker et al. and thus placed in subtribe Ratzeburgiinae rather than Rottboelliinae. Groups affected by these discrepancies are noted with an asterisk in Figure 5. Percent of species threatened ranged from 0-26.1% of each subtribe. The highest percentage is in Tripsacinae, a slightly misleading number since this subtribe also includes the largest percentage of species formally assessed by IUCN. The next highest percentages are Rhytachninae and Chionachninae, with 15.4 and 16.7% potentially threatened, respectively.

**Figure 5.**
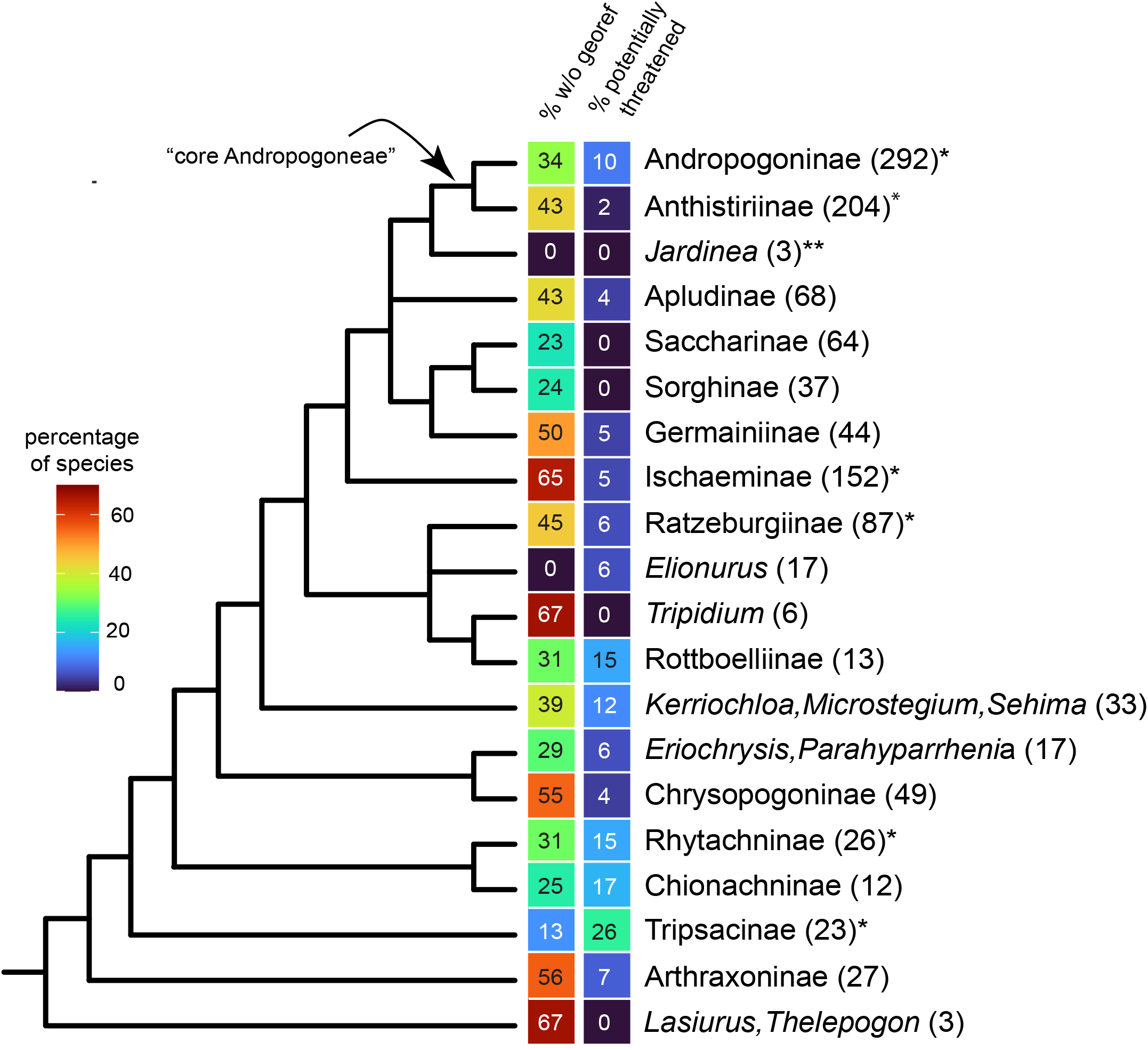
Phylogeny of subtribes of Andropogoneae, following Welker, McKain et al. (2021). Percentage of species with insufficient locality data for assessment (left gradient) and percentage potentially or actually threatened (right gradient). Numbers in parentheses = total number of species in the subtribe or clade, accordingly to Welker, McKain et al. (2021). *Subtribe contains threatened species that have been fully assessed for IUCN. \*\**Jardinea* species are placed in *Phacelurus* in the WCSP species list, but the two genera are unrelated.

The apparently low percentage of threatened species reflects the general paucity of locality data (Figure 5). In general, the percentage of species lacking digitized data is appreciably higher than the percentage that is potentially threatened. For example, while only 8 of the 152 species of subtribe Ischaeminae (5.3%) are potentially threatened, 99 (65.1%) lack enough locality data for even the very basic assessments that we have undertaken here.

### 3.4 Comparison to species with full assessments

Of the 100 Andropogoneae species already assessed for the Red List (Supplemental Table 1), 57 were also included in our analyses, which assigned the same extent of risk to 45 of them. ConR estimated a greater risk than the Red List for four species, and a lower risk for six. This was expected, as Red List assessments include information on population decline and location threats, neither of which is offered by rapid analyses. We did assess two species designated as Data Deficient (DD) in the Red List, suggesting that reassessment is warranted. We could not assess 42 of the Red List species because they lack digitized locality data; of those, 18 are designated as threatened. The Red List also assessed *Zea mays*, whereas we excluded it.

Combining the IUCN data with our other assessments, we suggest ∼11% of assessed Andropogoneae may be threatened.

### 3.5 Crop wild relatives

We addressed the conservation status of wild relatives of the major crops in Andropogoneae (maize, sorghum, sugarcane). Excluding *Zea mays, Sorghum halepense*, and *S. bicolor*, we filtered the species list for the 96 species in the genera *Zea, Tripsacum, Sorghum, Saccharum, Miscanthus, Hemisorghum*, and *Cleistachne*, which are the groups most likely to include crop wild relatives (Supplemental Table 3). The results mirrored those of the full data set: 26% of the species lacked sufficient records for assessment, 11.5% spanned the 180th meridian, and 62.5% could be assessed. Most of the latter group had EOO ratings of LC/NT; 10 species were estimated as either VU or EN under AOO, justifying further exploration.

The IUCN Red List has already assessed 27 species within the Andropogoneae crop wild relative genera. Of those, *Zea diploperennis, Tripsacum zopilotense, T. maizar, T. intermedium*, and *T. peruvianum* are EN, *Z. luxurians* is VU, and *Z. perennis* is CE (IUCN, 2020). All seven were Red Listed using criterion B, except *Z. perennis*, which was assessed with criterion A (population size reduction) (IUCN, 2020). Our analyses for the six species ConR could assess resulted in EOO ratings of LC/NT. None of the species newly assessed here were estimated to be threatened.

## 4 DISCUSSION

We show here that existing tools can provide rapid conservation assessments for a large set of species (1,127), dividing them quickly into those that are a) data deficient, b) likely not globally threatened, and c) in need of immediate attention. In Andropogoneae, over half the species lacked sufficient locality data in online databases, with five countries accounting for much of the missing data (Figure 4). These regions are clearly targets for enhanced collecting and digitization efforts. This percentage is similar to that reported by Zizka et al. (2021) for Orchidaceae, in which only 47% of the known species could be assessed. Insufficient geographic knowledge is also a pervasive problem among crop wild relatives (Khoury et al., 2020; Castañeda-Álvarez et al., 2016).

Of the species assessed, over 90% are likely not under imminent threat globally, although follow-up regional analyses and assessments may be warranted. Meanwhile, the most pressing need for further work is among the 50 species classified as potentially threatened by at least one assessment tool (Figure 6). We suggest that the next step is to develop a “sliding scale of concern” to consider acceptable thresholds for 1) extent of occurrences, 2) number of unique occurrences, 3) number of locations, and 4) regional species richness.

**Figure 6.**
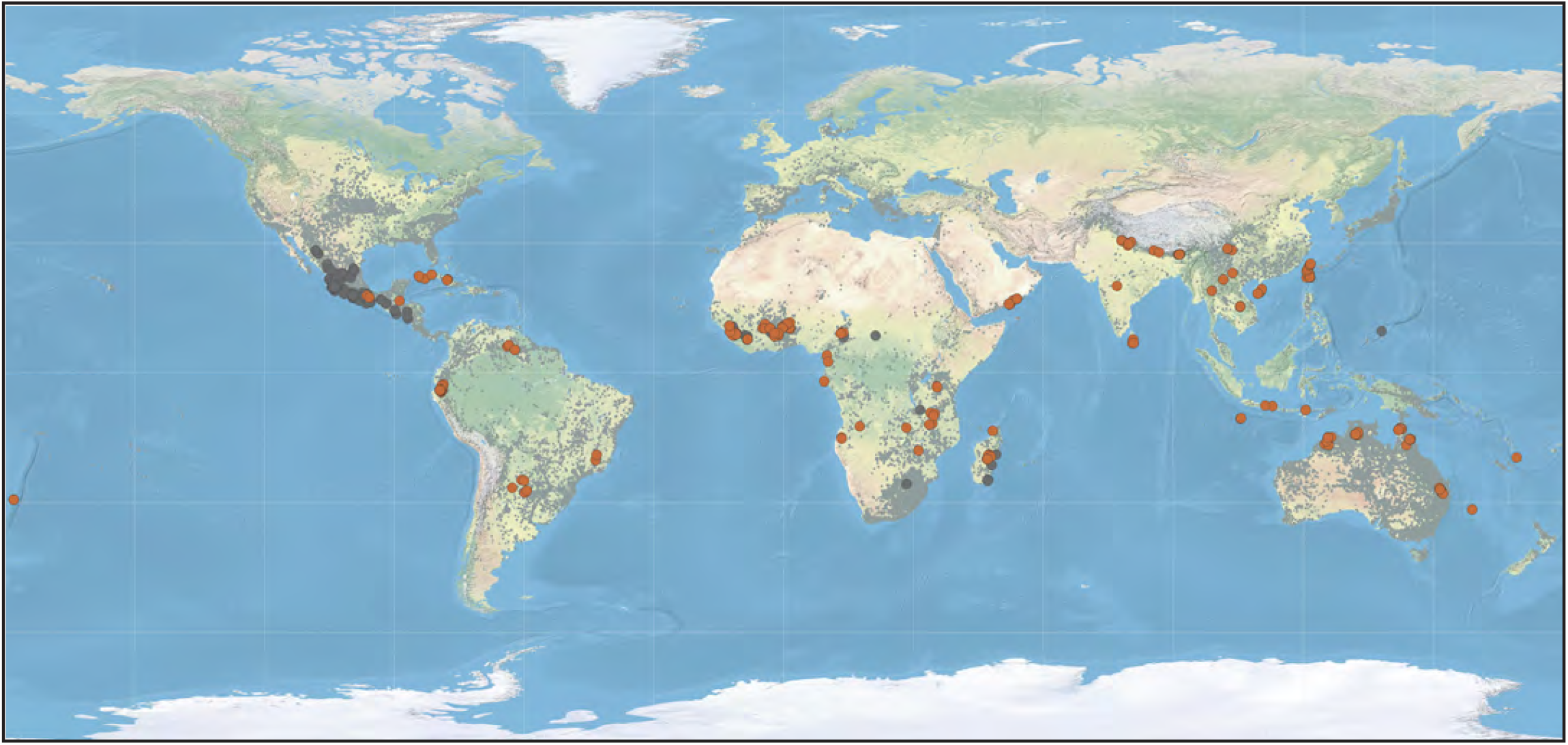
Global location of potentially threatened species (orange) and IUCN Red List-assessed taxa (dark gray) layered on estimated range of Andropogoneae (light gray). Mapped with QGIS-LTR Version 3.22.4. Graticules from Natural Earth.

Criterion B is only the first step toward a full Red List assessment, and assessments relying primarily on this criterion present clear limitations. Plant collection is often biased by taxon and locality (Walker et al., 2020; Lughadha et al., 2020). As seen in Figure 2E, sparse records at high latitudes of both hemispheres likely reflect climate preferences of most species whereas the lack of records in, for example, India, almost certainly reflects lack of collecting or digitization. Nonetheless, criterion B is often the necessary first step for assessments of plants (Schmidt et al., 2017; Pérez-Sarabia et al., 2020).

Assessment under Criterion B lends itself to automation (Zizka et al. (2021). We have found here that ConR produces EOO and AOO values that are virtually identical to those of GeoCAT, the tool that is approved by the IUCN for use in Red List entries, and it assigns most of the same taxa to the combined category LC/NT. The batch processing and automated output of ConR can assess hundreds of species at once and thus is useful for rapidly setting priorities. Although not attempted in the current study, it can also integrate data on protected areas (where available) and overlay them on EOO maps for further assessment of potential threats. ConR does fail when species distributions cross the 180th meridian, but this only affected a small percentage of species in our study. In the near future, we expect that ConR will be surpassed by newer, even more sophisticated tools using automation and deep learning such as IUC-NN (Zizka et al., 2021).

Phylogenetic diversity can also be used for setting conservation priorities (Li et al., 2017; Forest et al., 2007; Davies, 2019). Placing potentially threatened species on a phylogeny is becoming easier, with phylogenies are increasingly available for many plant clades. More elaborate analyses (e.g. Evolutionarily Distinct and Globally Endangered (EDGE) (Isaac et al., 2007) could then be done, although these go beyond the rapid triage that we are attempting here.

Reconciliation of names and synonyms was the most time-consuming step of the process used here. More robust tools for automated retrieval of currently accepted taxonomy will directly benefit rapid conservation assessment. Taxonomic irregularities are inevitable in any large clade and need to be addressed, particularly for apparently rare species that may be masquerading under misapplied (or simply misspelled) names. The TNRS (Boyle et al., 2013) is a big step forward toward automation, although the most recent update was released after we had largely completed the taxonomic component of this project. BIEN R itself clears up some taxonomic errors, but still requires additional checks for taxa that have multiple names in common use. We note that preliminary conservation assessment relies on good taxonomy but can proceed even while species limits are being reconsidered.

Among the crop wild relatives, the Red List listed six as threatened, whilst ConR suggested LC/NT. These seemingly contradictory results further support the argument that these preliminary assessments should serve as a first pass; while a taxon may be widespread, a closer regional analysis may identify specific threats like major changes in land use.

Obvious next steps for our priority species would be in-depth approaches like gap analysis (Sowa et al., 2007; Carver et al., 2021), connectivity analysis (Fajardo et al., 2014), or ecological modeling (Rodríguez et al., 2007), to name just a few. We hope to provide a basis for identifying and pursuing such studies, as our methods could be applied to many other groups.

Our work highlights three focal areas for conservation efforts: 1) Massively increased digitization and high-quality georeferencing of existing herbarium collections, particularly in species-rich regions; 2) Authoritative lists of species names and synonyms; and 3) Rapid automated tools for assessment of risk. Large-scale and preliminary conservation assessments deserve cautious evaluation, but are increasingly necessary to accelerate the process of predicting extinction risks of plants.

## Acknowledgements

We thank the Missouri Botanical Garden’s Burgund Bassuner, Roy Gereau, Patricia Barbera Sanchez, and Iván Jiménez for their initial guidance on IUCN Red Listing; and Maria Vorontsova at Royal Botanic Gardens Kew for access to GrassBase taxonomy data. This work was supported by NSF grant #1822330 to EAK.

## Author Contributions

TAE and EAK designed the research and wrote the manuscript. TAE, PM, and BB collected the data. TAE, PM, an EAK analyzed and interpreted the data. TAE and EAK created figures and maps.

## Data Availability Statement

Raw locality data, ConR maps for potentially threatened species, and R code are available on Github (https://github.com/ekellogg-lab/Androp_conservation)

## Supplemental material

**Supplemental Table S1**. Andropogoneae species assessed by the IUCN Red List as of 2021 with species name, Red List grade, and assessment year. LC = Least Concern, NT = Near Threatened, DD = Data Deficient, VU = Vulnerable, EN = Endangered, CR = Critically Endangered. *[A2][ac] where AC = “Population reduction observed, estimated, or inferred; where causes of reduction may not have ceased, may not be understood, or may not be reversible;” and ac = direct observation of decline in habitat quality, the area of occupancy (AOO), and/or extent of occurrence (EOO) (IUCN 2012). **D2 = Exclusive to VU category, where a time-sensitive “plausible future threat” is likely to cause a downgrade to CR or EX to already very small or restricted populations with <5 number of locations. B1 and B2 categories are summarized in Figure 1.

**Supplemental Table S2**. After taxonomic reconciliation, a final species list consisted of 1,130 names. *Sorghum bicolor, Sorghum halepense, and Zea mays* were excluded from conservation analysis but left in the species list. Species occurrences labeled with (†) denote estimated numbers, as ConR does not compute those that span the 180th meridian. (‡) denotes species that could not be computed because of insufficient data.

**Supplemental Table S3**. Congeneric taxa within crop wild relative genera *Cleistachne, Hemisorghum, Miscanthus, Saccharum, Sorghum, Tripsacum, Zea* were pulled from Supplemental Table S2 species list. Species occurrences labeled with (†) denote estimated numbers, as ConR does not compute those that span the 180th meridian. (‡) denotes species that could not be computed because of insufficient data.

